# Neural correlates of the effects of tDCS stimulation over the LIFG for phonological processing in dyslexia

**DOI:** 10.1101/522847

**Authors:** Lílian Rodrigues de Almeida, Peter C. Hansen

## Abstract

Findings of neuroimaging and brain stimulation research suggest that the motor system takes part in phonological processing at least to some extent in healthy speakers. Phonological processing involves a core network of brain regions, the dorsal pathway, where motoric aspects of speech sounds are analysed by the left inferior frontal gyrus (LIFG) and auditory aspects by the left superior temporal gyrus (LSTG). The extent to which each node of the dorsal pathway takes part in phonological processing has been shown to depend on the nature of the task and on the functional integrity of the network. Tasks of speech production rely more on the LIFG, and tasks of speech perception rely more on the LSTG. Persons with dyslexia (PWD) are known to present a deficit in phonological processing. Neuroimaging research has shown that dyslexia typically affects the LSTG, with hypoactivation, and the LIFG, with hyperactivation. Transcranial direct current stimulation (tDCS) has been widely used for cognitive research in humans. It has been recently suggested in the literature that perturbations induced by brain stimulation can cause the weights of nodes in cognitive networks to transiently rearrange. In this study we used tDCS and functional magnetic resonance imaging (fMRI) to investigate the functioning of the dorsal pathway for phonological processing in PWD with tasks of speech production and speech perception. We targeted the LIFG with anodal, cathodal and sham tDCS. For healthy speakers, cathodal tDCS should downregulate performance when the target had high relevance for the task, such as the LIFG for speech production. For targets of smaller relevance, improved performance should be observed due to compensation by the most relevant node(s). Anodal tDCS should improve performance as a function of the relevance of the target for the task. PWD were expected to deviate from this pattern to some extent, especially when compensation by the LSTG was needed during cathodal tDCS of the LIFG for a task of speech perception. Results corroborated the theoretical claim that codes for articulation take part in the processing of speech sounds. However, our findings showed that the PWD pattern of response to tDCS for phonological processing in tasks of speech production and speech perception differed from that expected for healthy speakers. Anodal tDCS of the LIFG induced larger facilitation for the speech perception than for the speech production task, as well as larger compensation for the latter under cathodal tDCS. Findings indicate that tDCS is a promising diagnosis tool for the investigation of alterations in phonological processing caused by dyslexia.

## 1 INTRODUCTION

In this study we investigated with functional magnetic resonance imaging (fMRI) the neural correlates of transcranial direct current stimulation (tDCS) of the left inferior frontal gyrus (LIFG) for phonological processing in persons with dyslexia (PWD). tDCS is a brain stimulation tool widely used for research in humans. tDCS modulates the excitability of the cerebral cortex, with anodal tDCS inducing improvement in performance and cathodal tDCS inducing decrease in performance, at least in the motor domain (Jacobson et al., 2012; Lang et al., 2004; Nitsche & Paul, 2000). In cognition, effects of tDCS on performance often deviate from this pattern (Jacobson et al., 2012). It has been suggested that the typical network structure of cortical brain regions underlying cognitive functions, as opposed to more circumscribed cortical regions underlying motor functions, would justify these differences (Jacobson et al., 2012). Findings in the literature suggest that the brain network nodes subserving a cognitive task differ in relevance, which can be rearranged to handle the task satisfactorily under endogenous or exogenous perturbations (Bestmann et al., 2008; Davis et al., 2008; Hartwigsen et al., 2012; Hartwigsen et al., 2013; Hartwigsen et al., 2016; Meinzer et al., 2009; Pirulli et al., 2014). Predictions for tDCS perturbations of cognitive networks are that anodal tDCS improves performance as a function of the relevance of the target for the task (or task load) (Bikson et al., 2013; Nozari, Woodard, & Thompson-Schill, 2014; Pope et al., 2015). Cathodal tDCS should decrease performance as a function of the relevance of the target for the task. However, if the target has low relevance, it may be that nodes of higher relevance can overperform to compensate the downregulation, resulting in improved performance or even a null result if compensation is not strong enough (Jacobson et al., 2012; Nozari, Woodard, & Thompson-Schill, 2014; Pirulli et al., 2014).

The neuroimaging and brain stimulation literature (e.g., Burton, 2001; Meister et al., 2007; Watkins & Paus, 2004) support the theoretical view of speech sound processing that considers that motor codes for articulation take part in speech perception at least to some extent (Liberman & Mattingly, 1985). The dorsal pathway for phonological processing, which connects the left inferior frontal gyrus (LIFG) to the left superior temporal gyrus (LSTG), has been widely investigated as the main brain network underlying phonological processing by integrating articulatory and acoustic information (Liebenthal et al., 2013; Saur et al., 2008) (Figure 1). The extent to which each node gets involved in a task would depend on the nature of the task and also on the integrity of the neural network. Tasks of speech production would rely more on the LIFG (Amunts et al., 1999; Eickhoff et al., 2009; Indefrey, 2011; Liakakis et al., 2011), whilst tasks of speech perception would rely more on the LSTG (Chang et al., 2010; Lee et al., 2012; Leonard & Chang, 2014; Liebenthal et al., 2013). However, since PWD have been shown to have an altered pattern of brain activity subserving phonological processing compared to healthy individuals (Brunswick et al., 1999; Carter et al., 2009; Georgiewa et al., 1999; Klingberg et al., 2000; Pagnotta et al., 2015; Rimrodt et al., 2010; Ruff et al., 2003; Waldie et al., 2013), predictions should be adjusted accordingly. The aim of this study was to investigate the effects of tDCS over the LIFG for phonological processing in tasks of speech production and speech perception in PWD, considering the particular altered pattern of brain activity for phonological processing presented by this population.

**Figure 1.**
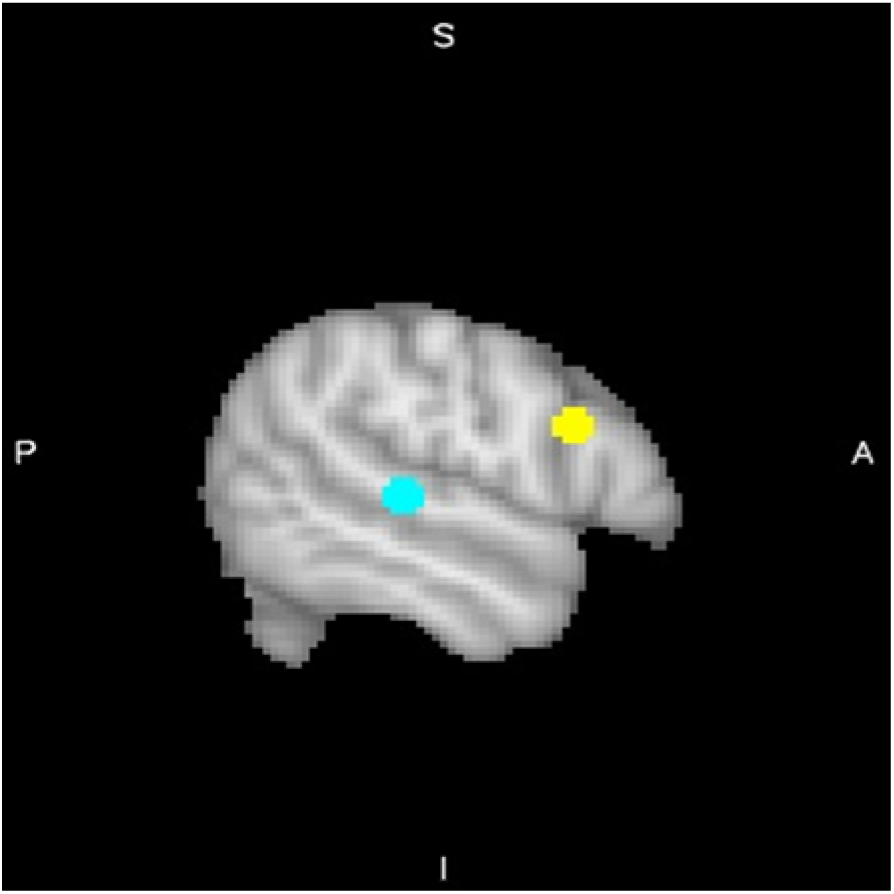
Dorsal pathway of phonological processing: LIFG (yellow) and LSTG (blue).

Developmental dyslexia is a hereditary language disorder characterised by a difficulty in learning to read that is not attributable to cognitive or sensory deficits or lack of educational opportunities (Ramus, 2004). It has been suggested that a deficit in phonological processing is always involved in all cases (Ramus, 2004), which is supported by neuroimaging studies that show altered patterns of brain activity and white matter integrity in areas involved with phonological processing (typically the LIFG and LSTG cortical areas and the arcuate and superior longitudinal fasciculi, Burton, 2001; Saur et al., 2008). The usual pattern of cortical alteration typically involves hypoactivation of the LSTG and hyperactivation of the LIFG (Brunswick et al., 1999; Georgiewa et al., 1999; Ruff et al., 2003). Communication between the frontal and temporal hubs has been shown to be weakened in both resting state and task-based analyses of functional connectivity (Schurz et al., 2015), as well as the relevant underlying white matter has been shown to have some degree of damage (Carter et al., 2009; Klingberg et al., 2000; Rimrodt et al., 2010). Furthermore, PWD have been demonstrated to have (maladaptive) compensatory mechanisms that commonly include the overactivation of the LIFG, RIFG (right inferior frontal gyrus) and RSTG (right superior temporal gyrus) (Pagnotta et al., 2015; Waldie et al., 2013). Costanzo et al.’s (2016a, 2016b) applied anodal tDCS to the LSTG and cathodal tDCS to the RSTG, and they found that brain stimulation was beneficial in improving the reading abilities in PWD. These results suggest that the LSTG has a potential to recover and that the RSTG might have a maladaptive role in dyslexia. Moreover, they support the view that tDCS shows promise for the treatment of dyslexia (Vicario & Nitsche, 2013).

In this study, compensatory LIFG and right cortical hyperactivation was expected to be evident for PWD. Predictions are presented in the subsections below by task. References to other more relevant nodes for compensation of the LIFG include the LSTG, the RIFG and the RSTG, i.e., the target network for this study. Although the LSTG seemed a preferred candidate among these network nodes because it is typically involved in phonological processing, the functional and structural connectivity between the LIFG and the LSTG in PWD could be impaired to some degree (Carter et al., 2009; Klingberg et al., 2000; Rimrodt et al., 2010; Schurz et al., 2015), and the alternative route of right compensation could prevail. Task difficulty could also potentially cause right hemispheric compensation (Gur et al., 2000), since the tasks used in this study were challenging in order to ensure participants and target engagement during brain stimulation.

### 1.1 Categorical perception

Findings in the literature support the assumption that categorical perception of speech has low task load for the LIFG and high task load for the LSTG (Chang et al., 2010; Lee et al., 2012; Leonard & Chang, 2014; Liebenthal et al., 2013; Rogers et al., 2014; Smalle et al., 2015; Watkins & Paus, 2004), at least for healthy young adults. This assumption should also apply for PWD. However, PWD are known to have a hyperactivated LIFG (Brunswick et al., 1999; Georgiewa et al., 1999; Hoeft et al., 2006; Ruff et al., 2003), that is likely to reflect maladaptive functioning (Brunswick et al., 1999; Meinzer et al., 2013). It was expected that anodal tDCS would increase the efficiency of the LIFG by reducing its level of activation (Antal et al., 2011; Meinzer et al., 2013). However, the effect could be considerably small, given the low task load involved. No significant changes from baseline were expected in network connectivity.

Cathodal tDCS was expected to increase the baseline level of neuronal activation (Antal et al., 2012) of the LIFG for PWD such as for healthy individuals, resulting in decreased efficiency of the LIFG to solve the task. Downregulation of a node of low relevance for the task was expected to induce compensation by more relevant network nodes to respond to the acute extra demand. In PWD, however, the LSTG is known to be hypoactivated (Brunswick et al., 1999; Georgiewa et al., 1999; Hoeft et al., 2006; Ruff et al., 2003). Results could therefore vary from successful compensation, but weaker than for healthy individuals, to an unsuccessful compensation. Strengthening of network connections should be observed for successful compensation.

### 1.2 Lexical decision

Lexical decision was assumed to have intermediate task load for the LIFG, because it involves reading, such as word naming, but not overt articulation. Anodal tDCS of the LIFG was therefore expected to induce facilitation, that should be higher than that expected for categorical perception. No significant changes from baseline were expected in network connectivity.

Similar to the predictions made for categorical perception, cathodal stimulation of the LIFG was expected to increase its level of neuronal activation (Antal et al., 2012) and decrease its efficiency in solving the task. Compensation of cathodal tDCS-induced downregulation of the LIFG by other network nodes was similarly expected through overactivation of network nodes or strengthening of their connections. However, compensation in lexical decision was likely to be more modest than in categorical perception, because the LIFG was assumed to have less room for compensation in the lexical decision task. As mentioned previously for the categorical perception task, compensation could be negatively affected by the LSTG hypoactivation present in PWD (Brunswick et al., 1999; Georgiewa et al., 1999; Hoeft et al., 2006; Ruff et al., 2003).

### 1.3 Word naming

The LIFG was assumed to be a node of high relevance for the word naming since this was a task placed on the production end of the speech perception to speech production range (Amunts et al., 1999; Eickhoff et al., 2009; Indefrey, 2011; Liakakis et al., 2011). As for healthy young adults, anodal tDCS of the LIFG would be expected to decrease the baseline level of activation (Antal et al., 2011) for PWD with consequent increased LIFG efficiency in solving the task. In PWD, anodal tDCS should decrease maladaptive hyperactivation, an effect expected to be beneficial (Meinzer et al., 2013). No significant changes from baseline were expected in network connectivity.

Similar to the predictions for healthy adults, cathodal tDCS was expected to increase the baseline level of activation of the target (Antal et al., 2012) in PWD with a consequent reduction in LIFG efficiency in solving the task. Compensation of cathodal tDCS-induced downregulation was expected to be unsatisfactory because the target was a node of high relevance for the task, and therefore, with less room for compensation.

### 1.4 Words and nonwords

The LIFG was assumed to be a node of higher relevance for nonwords than for words (Heim et al., 2005; Nosarti et al., 2010; Xiao et al., 2005). Anodal tDCS of the LIFG was expected to have for PWD the typical effect of reducing the target level of activation (Antal et al., 2011) with a consequent increased efficiency to solve the task. However, anodal tDCS effects were expected to be stronger for nonwords than for words, and results could vary from an apparent null effect of stimulation to a significant result. No significant changes from baseline were expected in network connectivity.

Cathodal tDCS of the LIFG was expected to increase the LIFG level of activation (Antal et al., 2012) and consequently decrease its efficiency in task solving also as a function of task load. Compensation of cathodal tDCS-induced downregulation by other network nodes was expected to be more successful for words than for nonwords because the LIFG was a node of comparatively less relevance for words. Compensatory activation of non-target nodes should be therefore higher for words than for nonwords, as well as increased the level of network connectivity. However, compensation in PWD was overall expected to be weaker than expected for healthy young adults because of the hypoactivation of the LSTG (Brunswick et al., 1999; Georgiewa et al., 1999; Hoeft et al., 2006; Ruff et al., 2003).

## 2 METHODS

### 2.1 Participants

Six right-handed (as assessed by the Annet’s (1972) handedness inventory) PWD who were native speakers of English were included in the sample (mean age: 20 years, SD: 1.94, 3 females). Twenty right-handed healthy young adults were used as controls for the assessment of reading abilities (mean age: 20.5 years, SD: 2.35, 9 females). Participants filled in safety questionnaires to unsure that they were eligible to undergo tDCS stimulation and MRI. All participants gave informed consent before taking part in the study, which was approved by the Central Ethics Committee of the University of Birmingham.

PWD in this study had a self-declared formal diagnosis of dyslexia. To confirm this diagnosis, PWD and controls undertook reading tasks (TIWRE and TOWRE, Torgesen et al., 1999; Reynolds & Kamphaus, 2007), and the cognitive tasks of block design, coding and picture completion from the WAIS IV battery (Weschler, 2008). PWD were expected to have reading difficulties, but similar performance to that of controls in other cognitive tasks (Ramus, 2004). Performance of PWD and controls on both task types confirmed the predictions. PWD had significantly lower reading performance than controls, particularly in the phonemic decoding task (accuracy: t(5.70) = −6.24, p < 0.001; reading speed: t(39) = 3.49, p < 0.01). No significant difference between PWD and controls was observed in the other cognitive tasks.

### 2.2 MATERIALS

#### 2.2.1 Tasks and stimuli

The categorical perception task involved the judgment of ten speech sound tokens from a continuum of synthesized speech between /ba/ and /da/, which should be categorised as either of the endpoints. The continuum was generated with a SenSyn Klatt synthesizer through the manipulation of the second and third formants of the endpoints. We used Raizada and Poldrack (2007) stimuli, who describe them in detail in their paper. The sound tokens had 300 ms and were repeatedly presented in randomised order for an unequal number of times each. The two more extreme tokens corresponding to the endpoints were presented 30% of the times, and the remaining tokens, which were more challenging to categorise, were presented 70% of the time. An MRI compatible headset (ConFon Electro Dynamic Headphones; MR confon GmbH, n.d.) was used to deliver the stimuli. Participants made their judgment by pressing the corresponding button with the left hand on a button box.

For the lexical decision and word naming tasks, an equal number of words and nonwords were randomly presented on the screen for 500 ms each per run. Stimuli were presented between two aligned vertical bars that stayed visible throughout the whole run. Different, but matched, lists of stimuli were used between the two runs, and were the same for both tasks.

Six letter words and nonwords were generated for the lexical decision and word naming tasks. Words were generated with the VWR R package (Keuleers, 2013) from a list of 66,330 English words from the CELEX lexical database (Baayen et al., 1995). Words were controlled for orthographic neighbourhood density (OLD20 score; Yarkoni et al., 2008) and frequency (SUBTLEXus database of word frequency for American English; Brysbaert & New, 2009). The summary statistics for word frequency (frequency count: frequency per million words) were: mean = 503.7, SD = 1966.4, min = 3 and max = 22040. Words generated with the VWR R package for which frequency was not found were excluded and replaced. Nonwords were generated with the Wuggy pseudowords generator (Keuleers et al., 2010). They were matched to the list of words by OLD20 score. The summary statistics for the OLD20 scores were mean = 1.88 and SD = 0.32 for words and mean = 1.88 and SD = 0.27 for nonwords. No significant difference between the OLD20 of words and nonwords was observed.

In the lexical decision task, participants should judge if the stimulus was a real word or a nonword by pressing the corresponding button with the left hand. In the word naming, all stimuli presented should be read aloud as promptly as possible. Voice responses were recorded with an MRI compatible microphone (Optoacoustics’ FOMRI III+ Noise Cancelling microphone; Optoacoustics Ltd., n.d.). However, it was not possible to filter the voice responses out of the scanner noise, and therefore the onset voice responses could not be used in the analyses.

We used a rhyme judgment task as a “warming up” task for the initial period of stimulation with tDCS to ensure enough time for tDCS to start to have an effect on behaviour (Nozari, Arnold, & Thompson-Schill, 2014) before the experimental tasks were presented. For the rhyme judgment task, pairs of words were randomly presented on the screen between aligned vertical bars for 900 ms each pair. Participants should judge each pair as a pair that rhymed or a pair that did not rhyme by pressing the corresponding button with the left hand. A single run of this task was presented between the two blocks of experimental tasks with an inter-trial interval of 2 s. Forty pairs of stimuli that rhymed and forty pairs of stimuli that did not rhyme were presented. Within each of these types of stimuli, half of them consisted of a pair of orthographic similar words and the other half consisted of a pair of orthographic dissimilar words. Stimuli consisted of a subset of McNorgan and Booth (2015)’s list of pairs of words.

All experimental tasks (categorical perception, lexical decision and word naming) had 60 stimuli per run and two runs per session (each session corresponded to one tDCS condition, i.e., anodal, cathodal or sham). Stimuli were presented with a variable inter-trial interval that followed a Poisson distribution whose mean was of 9.5 s. Visual stimuli were delivered with the Presentation software (version 18.3, Neurobehavioral Systems) via projector during the fMRI sessions.

#### 2.2.2 tDCS

Direct current with 2 mA of intensity was delivered with an MRI compatible neuroConn tDCS device (neuroConn GmbH, n.d.) through 5 x 5 rubber electrodes. With assistance of an EEG cap, the active electrode was placed on the LIFG, F5 according to the 10–20 international EEG system (Jasper, 1958), and the reference electrode was placed over the right supraorbicular region. Real tDCS conditions had a duration of stimulation of 20 minutes. The sham condition lasted 30 s, within the typical range of duration that is not enough to modulate brain function and therefore ensures a satisfactory placebo (Nitsche et al., 2008). For all conditions, stimulation started with the direct current increasing from zero to 2 mA with a ramp of 10 s and finished by decreasing the current from 2 mA to zero with a ramp down of 10 s. Ten20 conductive paste was applied to the electrodes to reduce scalp electrical resistance. This paste was used in lieu of the typical saline solution to avoid drying out of the electrodes during the experiments, since participants would be wearing them for a long time before the brain stimulation started.

### 2.3 PROCEDURE

The experiments were run with a within-subject design with a single blind protocol, where participants were unaware of the tDCS condition of each session. tDCS conditions of anodal, cathodal and sham were presented in different sessions and counterbalanced across participants.

Participants were given practice for the experimental tasks and for the rhyme judgment task prior to the experimental sessions. Written instructions were presented on the screen before each task, and orally reinforced by the researcher in charge of the session. Participants were reminded of responding to each task as appropriate as quickly as possible.

#### 2.3.1 Design of the experimental sessions

Task order presentation was counterbalanced across participants, but kept the same across the two blocks of a session and across all the sessions of each participant. Categorical perception, lexical decision and word naming were presented in two blocks, one for baseline and the other one under brain stimulation (online run). The rhyme judgment task was presented between blocks, in the beginning of the tDCS stimulation (see Figure 2). The effects of the current was assumed to be the same for all the experimental tasks of the online run, since the effects of the current has already been shown in the literature to persist for minutes after the end of the stimulation (Mangia et al., 2014). fMRI was acquired during the experimental tasks, with one scan per task. The rhyme judgment was run without scanning. A structural anatomical scan of each participant was acquired after the experimental runs of any one of the three tDCS sessions.

**Figure 2.**
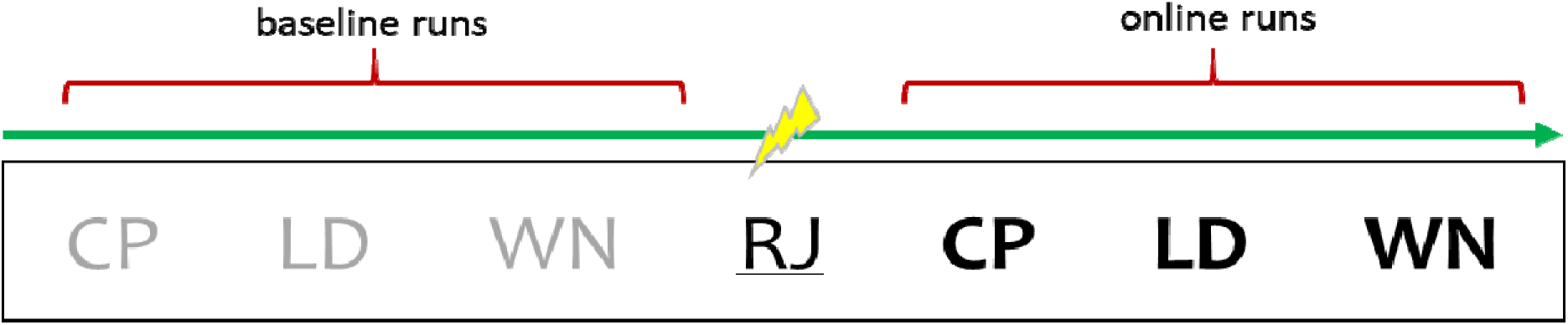
Schematic representation an experimental session. Experimental tasks (CP, lexical decision and word naming) are represented in grey for the baseline and in black for the online runs. Thunder ray symbol indicates the onset of the 20 minutes tDCS stimulation, which overlaps with the onset of the rhyme judgment task (RJ), underlined.

#### 2.3.2 MRI acquisition parameters

A 32 channel head-coil 3T Phillips Achieva scanner was used to collect MRI data at the Birmingham University Imaging Centre (BUIC). 240 T2*-weighted gradient-echo EPI volumes were acquired per scan (or experimental run), with a repetition time (TR) of 2.5 s, echo time (TE) of 34 ms, flip angle (FA) of 77°, slice thickness of 3 mm, voxel size of 3 mm^3^, field of view (FOV) of 240 x 130 x 240 mm and acquisition matrix of 80 x 80. Each EPI volume had 43 axial oblique slices, that was enough to cover the whole cortex. Slices were acquired in sequential descending order. There is some recommendation in the literature for the use of sparse sampling for speech production and speech perception tasks in order to minimise motion artifacts caused by articulation and the interference of background scanner noise with the reception of speech (Raizada & Poldrack, 2007; Ulm et al., 2015). However, this would bring the caveat of sequence variability between tasks, as well as diminished statistical power, due to the reduction of number of stimuli to fit the task within the same duration. To avoid these issues, the typical non-sparse sampling sequence was equally used for all the tasks. A pilot study was run to ensure the feasibility of the experimental tasks with the typical sequence. The structural anatomical scan was an isotropic T1-weighted gradient echo image with the following parameters: 175 sagittal slices, TR = 8.4 ms, TE = 3.8 ms, flip angle of 7° and voxel size of 1 mm^3^.

### 2.4 ANALYSES

#### 2.4.1 Preprocessing

The FMRIB Software Library (FSL; Jenkinson et al., 2012; Smith et al., 2004; Woolrich et al., 2009) was used for preprocessing of functional and structural images, and to analyse the fMRI data. Non-brain tissue was removed from structural anatomical images (T1) with the FSL BET (v.2.1) tool (Smith, 2002). Functional images received regular-down slice timing correction. Images were spatially smoothed with a Gaussian kernel of 4.5 mm (1.5 times one dimension of the isotropic 3 mm^3^ voxel). Motion correction of the functional data was performed by using the MCFLIRT tool (Jenkinson et al., 2002) and the ICA-AROMA (ICA-based Automatic Removal Of Motion Artifacts) tool (Pruim, Mennes, Buitelaar et al., 2015; Pruim, Mennes, van Rooij et al., 2015). The MCFLIRT applied rigid body transformation with the middle image as reference. The ICA-AROMA was used to identify and remove motion-related ICA components from the data. Temporal filtering was applied after ICA-AROMA motion correction. The high pass Gaussian weighted filter cut-off was of 50 s.

Multi-stage registration was performed. A 6 DOF affine registration was used to register functional images to individual anatomical space with the FSL FLIRT tool (v.6.0) (Jenkinson & Smith, 2001; Jenkinson et al., 2002). A non-linear registration (warp resolution 10 mm) of each functional image into standard MNI space was then performed with the FSL FNIRT tool (Andersson et al., 2007a, 2007b).

#### 2.4.2 Data analyses

##### 2.4.2.1 Whole brain analyses

Whole brain analyses were conducted to investigate the overall brain activation induced by the factors of task, tDCS and population (data not reported). They were conducted as a first stage of analyses before mean activation of regions of interest, i.e., regions of the target network for phonological processing (LIFG, LSTG, RIFG and RSTG), could be calculated.

Data were analysed with the FSL FEAT v.6.0 tool (Woolrich et al., 2001; Woolrich et al., 2004). A general linear model (GLM) with local autocorrelation correction (using FILM prewhitening; Woolrich et al., 2001) was used to analyse all conditions at the individual level. Each of the functional runs in a session corresponded to one task (either categorical perception, lexical decision or word naming), one tDCS condition (either anodal, cathodal or sham for a particular session) and one repeat (either baseline or online). In the first level of analysis, only task was therefore modelled as factor of interest for each run.

For the lexical decision and the word naming tasks, different stimulus types were entered into the design matrix as separate covariates, i.e., words and nonwords were modelled separately. Onset of responses were included in the design matrix as nuisance covariates whenever available (not available for word naming). Stimuli presentation and responses had their onset and duration (as described in section 2.2.1 Tasks and stimuli) modelled. Button responses were given a notional duration of 100 ms. Time courses associated with each event where onset and duration were modelled were convolved with a double-gamma HRF (Hemodynamic Response Function). Temporal filtering was applied and temporal derivatives were added to the model as separate nuisance covariates in order to improve the model fit. Motion parameters generated by MCFLIRT were also included as nuisance covariates to regress out unwanted influence of motion on performance (Johnstone et al., 2006). T-contrasts were generated for the mean of all stimulus types versus rest for all the tasks. In addition, for the lexical decision and word naming tasks, t-contrasts were generated for the mean of each stimulus type, i.e., words and nonwords, versus rest.

Second level analyses were carried out with fixed effect models by participant with the contrast images from the first level analyses as input, i.e., contrast images for the mean across all the stimuli of each task (and mean across stimulus types for lexical decision and word naming) per run (baseline and online) and tDCS condition (anodal, cathodal and sham). Each task (and stimulus type for lexical decision and word naming)/tDCS combination was set up separately as a covariate of interest in the design matrix. The difference between the online and the baseline repeat was set up in the design matrix within the covariates for task/tDCS combinations. T-contrasts were set up to perform the differences between real tDCS (anodal or cathodal) and sham for each task (and stimulus type for lexical decision and word naming).

Group analyses were carried out with random effect models using FLAME stage 1 (Beckmann et al., 2003; Woolrich et al., 2004; Woolrich, 2008). Gaussian Random Field Theory thresholding was applied to the statistical maps, with a value of Z > 2.3 at the voxel level and p < 0.05 at the cluster level, corrected for multiple comparisons. Z-value activation maps were produced for each contrast. The second level output images of each participant entered the models as input. Mean t-contrasts (one-sample t tests) were set up to analyse the group mean brain activation for each task (and stimulus type for LD and WN)/tDCS combination from the second levels.

##### 2.4.2.2 ROI analyses

ROI analyses were conducted to investigate patterns of activation in the target network (which consisted of the LIFG, the LSTG, the RIFG and the RSTG) under tDCS stimulation.

###### 2.4.2.2.1 ROI definitions and ROI-based data measurements

The network of interest for the fMRI experiments consisted of the two typical nodes involved in phonological processing, LIFG and LSTG (Burton, 2001; Liebenthal et al., 2013; Saur, 2008), and their right homologues, RIFG and RSTG. Four corresponding ROIs were created with FSL command line tools as a 6 mm radius sphere centred at coordinates of interest in MNI space.

Coordinates for LIFG and LSTG were obtained from meta-analyses of functional brain activation associated with the CP, lexical decision and word naming tasks. These were carried out with the Neurosynth software and database (Neurosynth, 2018; Yarkoni et al., 2011a, 2011b). The search for each task used, respectively, the keywords “speech perception”, “lexical decision” and “speech production”, and yielded three forward inference statistical maps Benjamini-Hochberg corrected for multiple comparisons with a threshold of 0.01 (see the Neurosynth website and Yarkoni et al., 2011a, 2011b references for further information). By using FSL command line tools, the intersection between the three statistical images was obtained. The resulting image was submitted to a cluster analysis with a Z threshold of 2.3. Clusters corresponding to the LIFG and the LSTG in the Harvard-Oxford cortical atlas available in FSL (Desikan et al., 2006; Frazier et al., 2005; Goldstein et al., 2007; Makris et al., 2006) were identified through their centre of gravity (COG), that is an average of the coordinates within the cluster weighted by intensity. These were then chose as the coordinates for LIFG and LSTG. Coordinates for the right homologues RIFG and RSTG were the same as those for the left ROIs, but with the sign for the x coordinate reversed. The MNI coordinates for the four ROIs were x = −50, y = 14 and z = 24 (LIFG), x = −58, y = −28 and z = 4 (LSTG), x = 50, y = 14 and z = 24 (RIFG) and x = 58, y = −28 and z = 4 (RSTG).

For each participant, mean percentage signal changes were obtained for each condition of interest per ROI with the FSL Featquery tool, based on whole brain analysis contrasts. Conditions of interest were the effect of task (and stimulus type for lexical decision and word naming) in combination with tDCS (second level contrasts) on brain activation. Contrasts involving the factor tDCS were defined with run 1 (baseline) subtracted from run 2 (online stimulation) and sham subtracted from real tDCS conditions (henceforth “anodal tDCS” or “cathodal tDCS”).

###### 2.4.2.2.2 Regression: effects of task and tDCS on mean brain activation per ROI

ROI mean activation measurements were fed into mixed effect linear regressions performed in R version 3.4.2 (R Core Team, 2017) to investigate, whenever applicable, the main effects of the within-subject factors of task, stimulus type (for lexical decision and word naming), tDCS and ROI on BOLD signal change, with participants included in the models as random effects. Post-hoc analyses were further set up as appropriate with contrasts to investigate whether each task (and stimulus type for lexical decision and word naming) in combination with tDCS was significantly different from zero per ROI. All contrast analyses were corrected for multiple comparisons using the Benjamini-Hochberg method (Benjamini & Hochberg, 1995).

###### 2.4.2.2.3 Partial correlation: ROI-based connectivity analyses per task and tDCS combination

Correlational analyses can provide indirect measurement of functional connectivity and have been extensively used for individual level analyses (e.g., Marrelec et al., 2006; Ryali et al., 2012; Sandberg, 2017), especially when precise prior information (e.g. temporal) for the connections between pairs of nodes, usually required to perform effective connectivity analyses, is not available. Partial correlation is therefore deemed to be a reasonable option (Marrelec et al., 2006). Furthermore, as connectivity analyses in this study were based on a previously defined network, partial correlation was considered more adequate than seed-based analyses, which are rather exploratory. For these reasons, partial correlation was the analysis of choice to investigate functional connectivity in this study.

Partial correlations using Pearson’s r, and their level of significance, were calculated for datasets of ROI mean brain activations with the PPCOR R package (Kim, 2015). These analyses were performed to investigate the relationships between each pair of nodes of the target network for the different conditions of task and tDCS to show the most prominent brain activity subserving performance. Datasets for each condition were selected according to contrasts of task or stimulus type (for lexical decision and word naming) in combination with tDCS.

## 3 RESULTS

Results of ROI analyses are presented in this section. Effects of task, stimulus type (for lexical decision and word naming), tDCS condition and ROIs on brain activation were analysed with mixed effect models for PWD. Relevant connections between ROIs were investigated with partial correlation analyses per task, stimulus type (for lexical decision and word naming) and tDCS condition. Both types of analyses are presented by task.

### 3.1 Categorical perception

#### 3.1.1 Task and tDCS effects on ROI mean brain activation

A 2 x 4 (tDCS x ROI) linear mixed effect model was fitted to the mean parameter estimates of ROI activation data of both anodal and cathodal tDCS conditions. No significant interaction between tDCS and ROI was observed.

Post hoc contrast analyses (Benjamini-Hochberg corrected for multiple comparisons) were performed, but none of them appeared significantly different from zero (Figure 3 and Table 1).

**Table 1.** Contrast analyses for fitted brain activations per ROI and tDCS

| Contrast | Estimate | SE | df | *t* | *p* |
| --- | --- | --- | --- | --- | --- |
| LIFG: anodal | -0.05 | 0.10 | 5 | -0.50 | 0.85 |
| LIFG: cathodal | -0.17 | 0.10 | 5 | -1.75 | 0.56 |
| LSTG: anodal | 0.03 | 0.10 | 5 | 0.28 | 0.85 |
| LSTG: cathodal | -0.07 | 0.10 | 5 | -0.69 | 0.85 |
| RIFG: anodal | -0.14 | 0.10 | 5 | -1.44 | 0.56 |
| RIFG: cathodal | -0.25 | 0.10 | 5 | -2.56 | 0.41 |
| RSTG: anodal | -0.06 | 0.10 | 5 | -0.64 | 0.85 |
| RSTG: cathodal | -0.02 | 0.10 | 5 | -0.21 | 0.85 |

**Figure 3.**
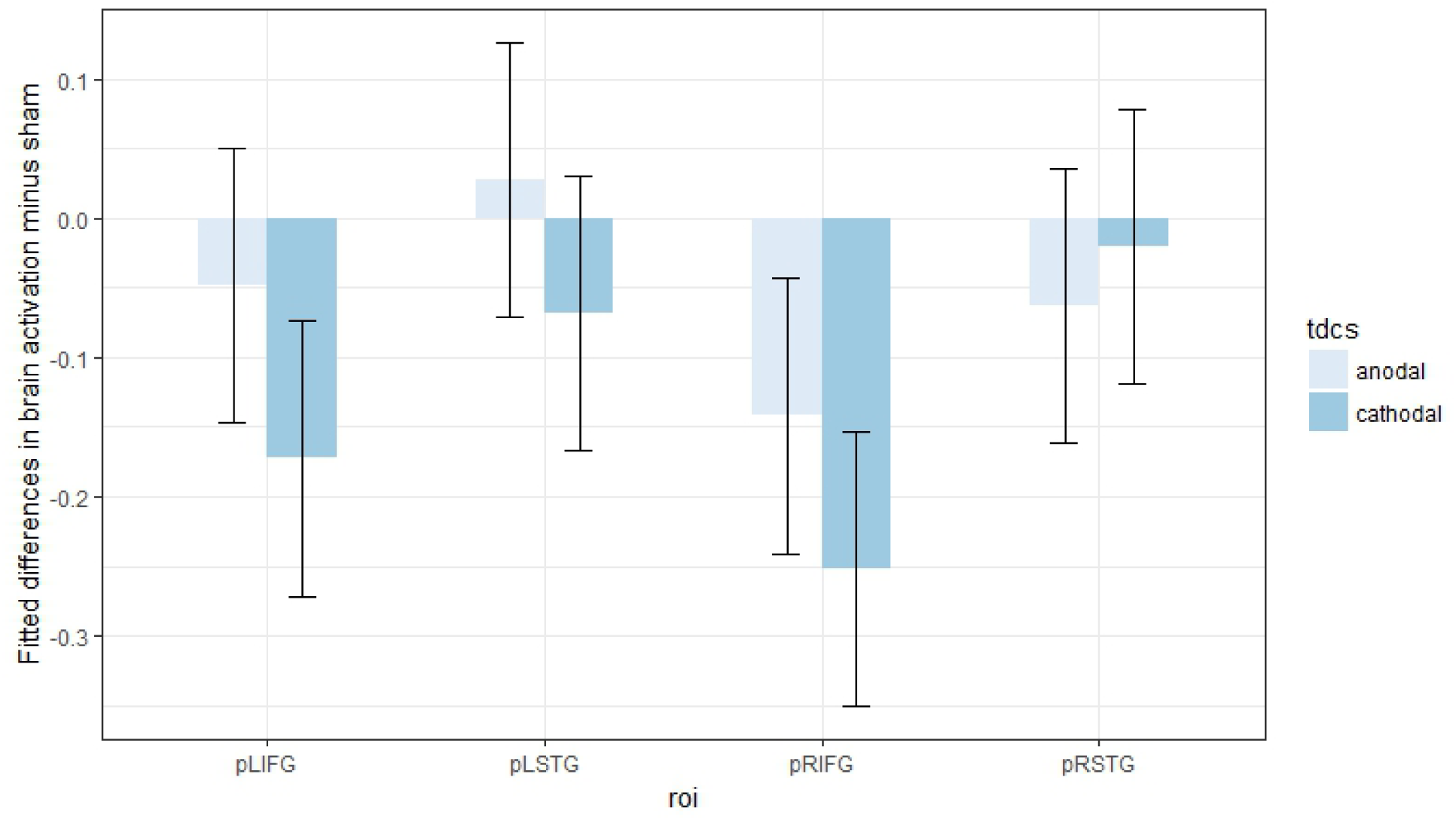
Fitted mean brain activation per ROI and tDCS for categorical perception. The x-axis displays the ROIs. The y-axis displays the fitted mean brain activation. Error bars represent the contrast estimate ± the pooled standard error.

#### 3.1.2 Connectivity analysis per tDCS condition

Partial correlation analyses were performed between the fitted mean brain activations of the target network ROIs by tDCS condition. Only anodal tDCS induced significant correlations, and these were: LSTG/RSTG and RIFG/RSTG (Tables 2 and 3).

### 3.2 Lexical decision

#### 3.2.1 Task and tDCS effects on ROI mean brain activation

A 2 x 4 (tDCS x ROI) linear mixed effect model was fitted to the mean parameter estimates of ROI activation data of both anodal and cathodal tDCS conditions. No significant interaction of tDCS and ROI was observed.

**Table 2.**
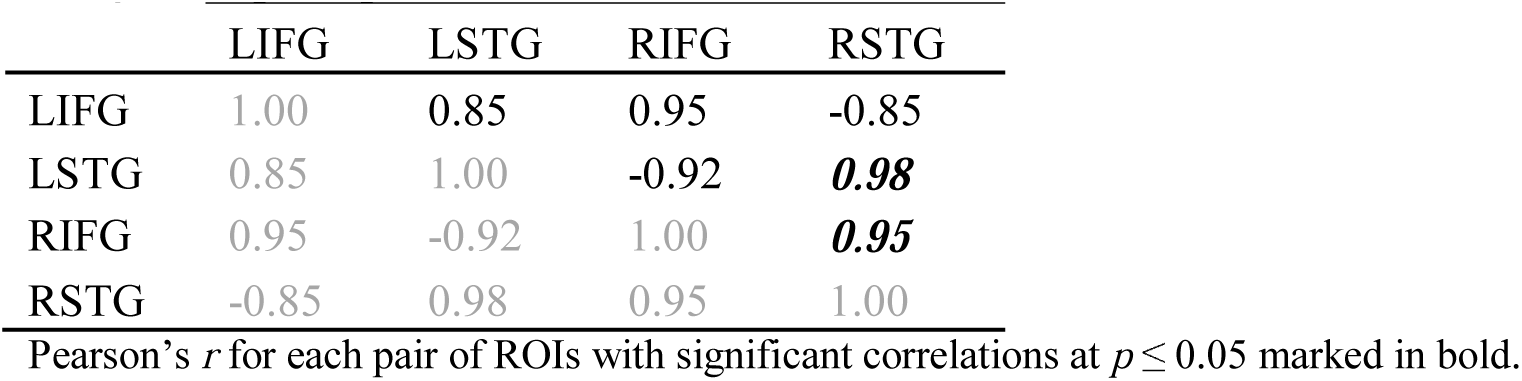
Partial correlation analyses for fitted mean brain activations under anodal tDCS in categorical perception

|  | LIFG | LSTG | RIFG | RSTG |
| --- | --- | --- | --- | --- |
| LIFG | 1.00 | 0.85 | 0.95 | -0.85 |
| LSTG | 0.85 | 1.00 | -0.92 | **0.98** |
| RIFG | 0.95 | -0.92 | 1.00 | **0.95** |
| RSTG | -0.85 | 0.98 | 0.95 | 1.00 |

Pearson's *r* for each pair of ROIs with significant correlations at $p \leq 0.05$ marked in bold.

**Table 3.** Partial correlation analyses for fitted mean brain activations under cathodal tDCS in categorical perception

|  | LIFG | LSTG | RIFG | RSTG |
| --- | --- | --- | --- | --- |
| LIFG | 1.00 | 0.55 | 0.78 | -0.51 |
| LSTG | 0.55 | 1.00 | -0.86 | 0.95 |
| RIFG | 0.78 | -0.86 | 1.00 | 0.89 |
| RSTG | -0.51 | 0.95 | 0.89 | 1.00 |

Pearson's *r* for each pair of ROIs. No significant correlation at $p \leq 0.05$ was observed.

Post hoc contrast analyses (Benjamini-Hochberg corrected for multiple comparisons) were performed, but none of them appeared significantly different from zero (Figure 4 and Table 4).

**Table 4.** Contrast analyses for fitted brain activations per ROI and tDCS

| Contrast | Estimate | SE | df | *t* | *p* |
| --- | --- | --- | --- | --- | --- |
| LIFG: anodal | 0.09 | 0.09 | 5 | 1.04 | 0.48 |
| LIFG: cathodal | 0.09 | 0.09 | 5 | 1.00 | 0.48 |
| LSTG: anodal | 0.02 | 0.09 | 5 | 0.23 | 0.83 |
| LSTG: cathodal | 0.02 | 0.09 | 5 | 0.28 | 0.83 |
| RIFG: anodal | 0.18 | 0.09 | 5 | 2.03 | 0.39 |
| RIFG: cathodal | 0.24 | 0.09 | 5 | 2.74 | 0.33 |
| RSTG: anodal | 0.13 | 0.09 | 5 | 1.51 | 0.48 |
| RSTG: cathodal | 0.11 | 0.09 | 5 | 1.26 | 0.48 |

**Figure 4.**
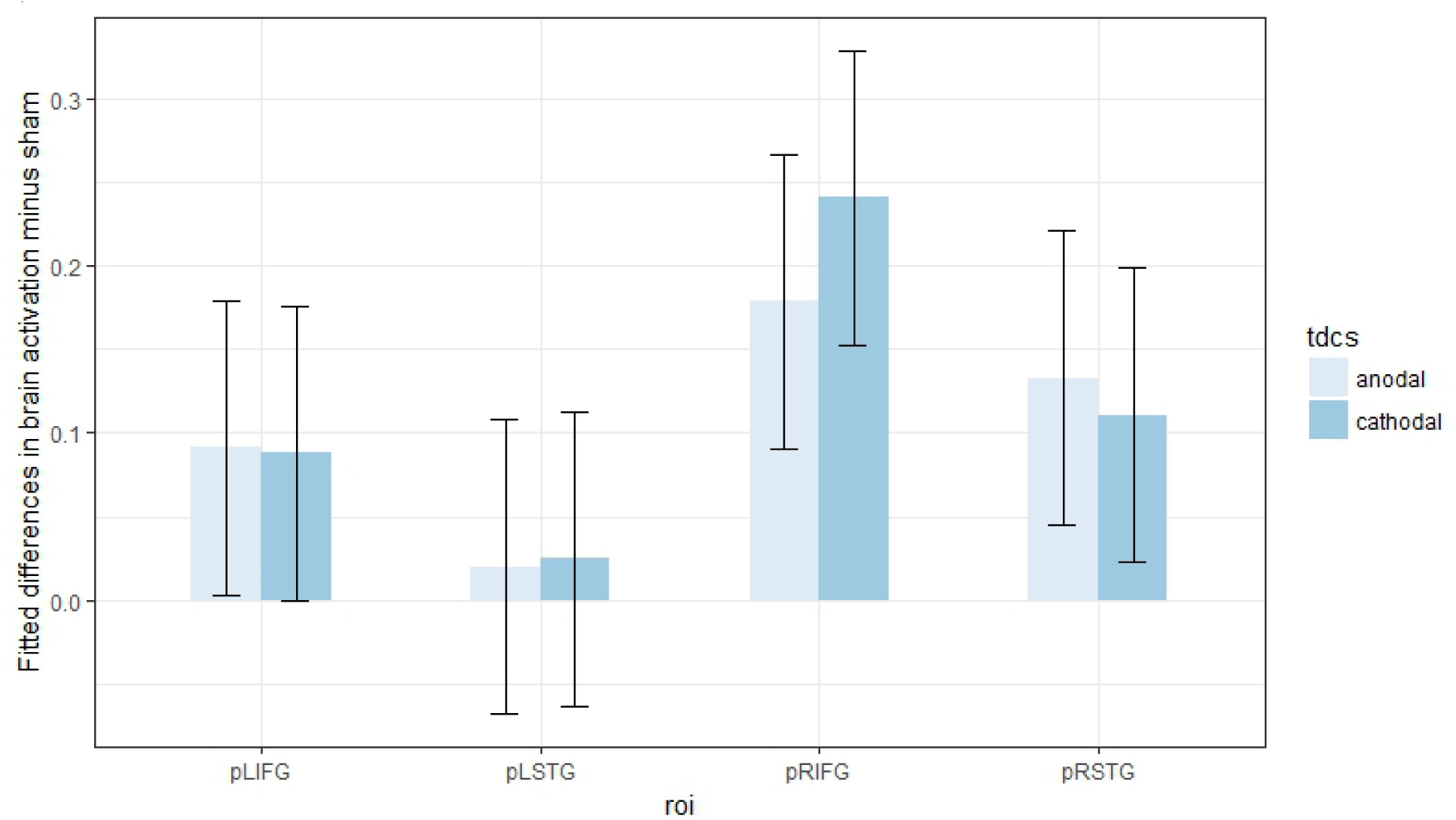
Fitted brain activation per ROI and tDCS for lexical decision. The x-axis displays the ROIs. The y-axis displays fitted mean brain activation. Error bars represent the contrast estimate ± the pooled standard error.

#### 3.2.2 Connectivity analysis per tDCS condition

Partial correlation analyses were performed between the fitted mean brain activations of the target network ROIs by tDCS condition. Neither anodal tDCS nor cathodal tDCS induced significant correlations at p ≤ 0.05 (Tables 5 and 6).

### 3.3 Word naming

#### 3.3.1 Task and tDCS effects on ROI mean brain activation

A 2 x 4 (tDCS x ROI) linear mixed effect model was fitted to the mean parameter estimates of ROI activation data of both anodal and cathodal tDCS conditions. No significant interactions were observed.

Post hoc contrast analyses (Benjamini-Hochberg corrected for multiple comparisons) were performed, but none of them appeared significantly different from zero (Figure 5 and Table 7).

**Table 5.** Partial correlation analyses for fitted mean brain activations under anodal tDCS in lexical decision

|  | LIFG | LSTG | RIFG | RSTG |
| --- | --- | --- | --- | --- |
| LIFG | 1.00 | -0.08 | 0.61 | 0.84 |
| LSTG | -0.08 | 1.00 | 0.50 | 0.48 |
| RIFG | 0.61 | 0.50 | 1.00 | -0.74 |
| RSTG | 0.84 | 0.48 | -0.74 | 1.00 |

Pearson's $r$ for each pair of ROIs. No significant correlation at $p \leq 0.05$ was observed.

**Table 6.** Partial correlation analyses for fitted mean brain activations under cathodal tDCS in lexical decision

|  | LIFG | LSTG | RIFG | RSTG |
| --- | --- | --- | --- | --- |
| LIFG | 1.00 | 0.01 | 0.30 | 0.20 |
| LSTG | 0.01 | 1.00 | -0.18 | 0.67 |
| RIFG | 0.30 | -0.18 | 1.00 | 0.60 |
| RSTG | 0.20 | 0.67 | 0.60 | 1.00 |

Pearson's $r$ for each pair of ROIs. No significant correlation at $p \leq 0.05$ was observed.

**Table 7.** Contrast analyses for fitted brain activations per ROI and tDCS

| Contrast | Estimate | SE | df | *t* | *p* |
| --- | --- | --- | --- | --- | --- |
| LIFG: anodal | -0.03 | 0.08 | 5 | -0.33 | 0.78 |
| LIFG: cathodal | 0.04 | 0.08 | 5 | 0.45 | 0.78 |
| LSTG: anodal | 0.05 | 0.08 | 5 | 0.55 | 0.78 |
| LSTG: cathodal | 0.04 | 0.08 | 5 | 0.48 | 0.78 |
| RIFG: anodal | -0.03 | 0.08 | 5 | -0.35 | 0.78 |
| RIFG: cathodal | 0.02 | 0.08 | 5 | 0.29 | 0.78 |
| RSTG: anodal | -0.04 | 0.08 | 5 | -0.42 | 0.78 |
| RSTG: cathodal | 0.07 | 0.08 | 5 | 0.87 | 0.78 |

**Figure 5.**
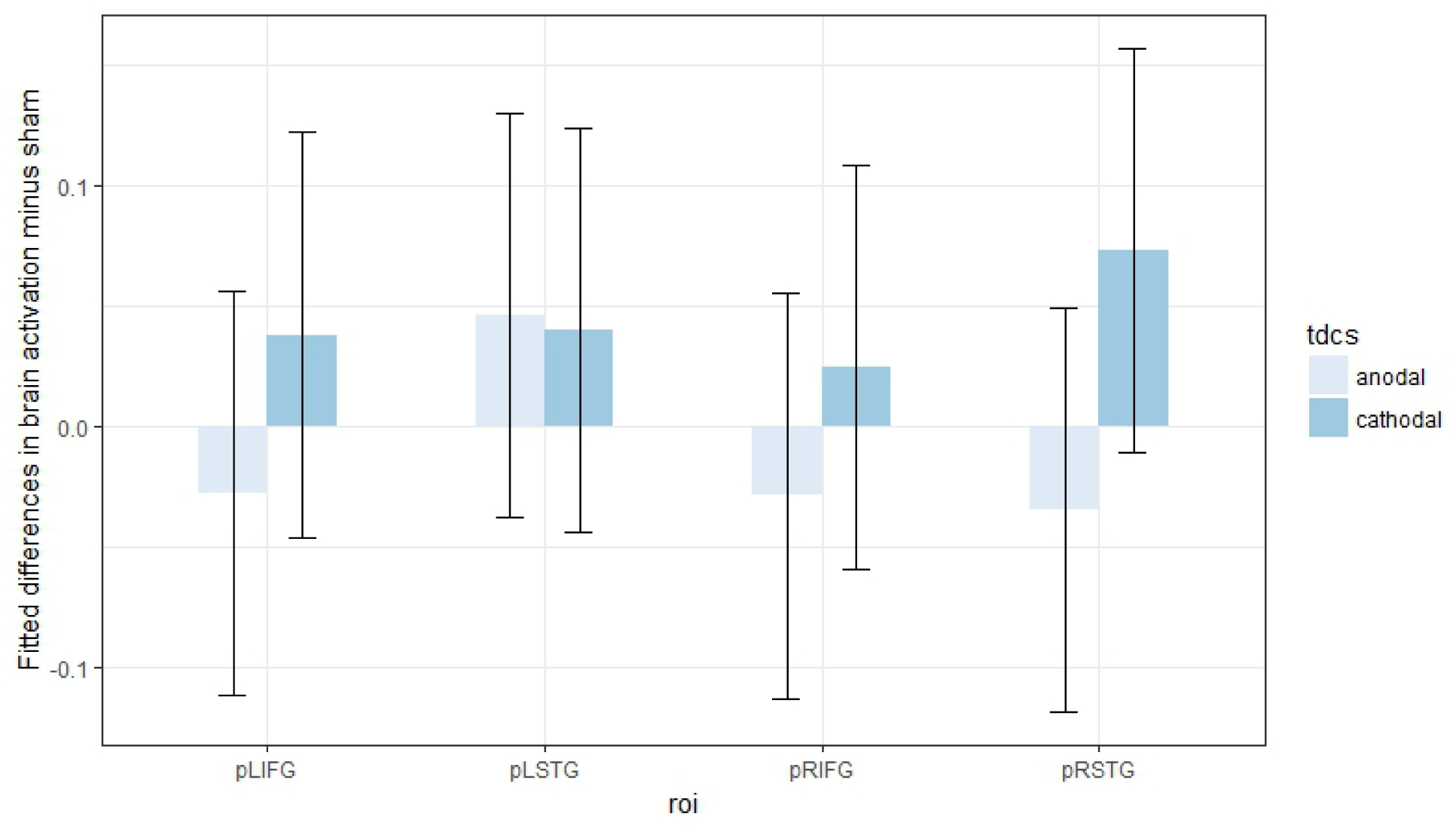
Fitted mean brain activation per ROI and task for word naming. The x-axis displays the ROIs. The y-axis displays the fitted mean brain activation. Error bars represent the contrast estimate ± the pooled standard error.

#### 3.3.2 Connectivity analysis per tDCS condition

Partial correlation analyses were performed between the fitted mean brain activations of the target network ROIs by tDCS condition. Anodal tDCS induced the frontal LIFG/RIFG significant correlation. Cathodal tDCS induced the left lateralised LIFG/LSTG significant correlation (Tables 8 and 9).

**Table 8.** Partial correlation analyses for fitted mean brain activations under anodal tDCS in word naming

|  | LIFG | LSTG | RIFG | RSTG |
| --- | --- | --- | --- | --- |
| LIFG | 1.00 | -0.10 | **0.96** | -0.94 |
| LSTG | -0.10 | 1.00 | 0.33 | -0.06 |
| RIFG | 0.96 | 0.33 | 1.00 | 0.94 |
| RSTG | -0.94 | -0.06 | 0.94 | 1.00 |

Pearson's $r$ for each pair of ROIs with significant correlations at $p \leq 0.05$ marked in bold.

**Table 9.** Partial correlation analyses for fitted mean brain activations under cathodal tDCS in word naming

|  | LIFG | LSTG | RIFG | RSTG |
| --- | --- | --- | --- | --- |
| LIFG | 1.00 | **0.98** | 0.67 | -0.18 |
| LSTG | 0.98 | 1.00 | -0.68 | 0.30 |
| RIFG | 0.67 | -0.68 | 1.00 | 0.30 |
| RSTG | -0.18 | 0.30 | 0.30 | 1.00 |

Pearson's $r$ for each pair of ROIs with significant correlations at $p \leq 0.05$ marked in bold.

### 3.4 Analysis of words and nonwords in lexical decision

#### 3.4.1 Stimulus type and tDCS effects on ROI mean brain activation

A 2 x 3 x 4 (stimulus type x tDCS x ROI) linear mixed effect model was fitted to the mean parameter estimates of ROI activation data of words and nonwords from the lexical decision task data. No significant interactions were observed.

Post hoc contrast analyses (Benjamini-Hochberg corrected for multiple comparisons) were conducted, but no significant result was observed (Figure 6 and Table 10).

**Table 10.** Contrast analyses for fitted brain activations per ROI and stimulus type

| Contrast | Estimate | SE | df | *t* | *p* |
| --- | --- | --- | --- | --- | --- |
| LIFG: nonword/anodal | 0.05 | 0.08 | 5 | 0.63 | 0.69 |
| LIFG: word/anodal | 0.10 | 0.08 | 5 | 1.29 | 0.48 |
| LIFG: nonword/cathodal | 0.02 | 0.08 | 5 | 0.24 | 0.86 |
| LIFG: word/cathodal | 0.11 | 0.08 | 5 | 1.42 | 0.48 |
| LSTG: nonword/anodal | -0.01 | 0.08 | 5 | -0.19 | 0.86 |
| LSTG: word/anodal | 0.05 | 0.08 | 5 | 0.65 | 0.69 |
| LSTG: nonword/cathodal | -0.03 | 0.08 | 5 | -0.41 | 0.80 |
| LSTG: word/cathodal | 0.07 | 0.08 | 5 | 0.94 | 0.63 |
| RIFG: nonword/anodal | 0.11 | 0.08 | 5 | 1.39 | 0.48 |
| RIFG: word/anodal | 0.21 | 0.08 | 5 | 2.72 | 0.33 |
| RIFG: nonword/cathodal | 0.14 | 0.08 | 5 | 1.74 | 0.48 |
| RIFG: word/cathodal | 0.28 | 0.08 | 5 | 3.58 | 0.25 |
| RSTG: nonword/anodal | 0.10 | 0.08 | 5 | 1.24 | 0.48 |
| RSTG: word/anodal | 0.13 | 0.08 | 5 | 1.63 | 0.48 |
| RSTG: nonword/cathodal | 0.06 | 0.08 | 5 | 0.76 | 0.69 |
| RSTG: word/cathodal | 0.13 | 0.08 | 5 | 1.67 | 0.48 |

**Figure 6.**
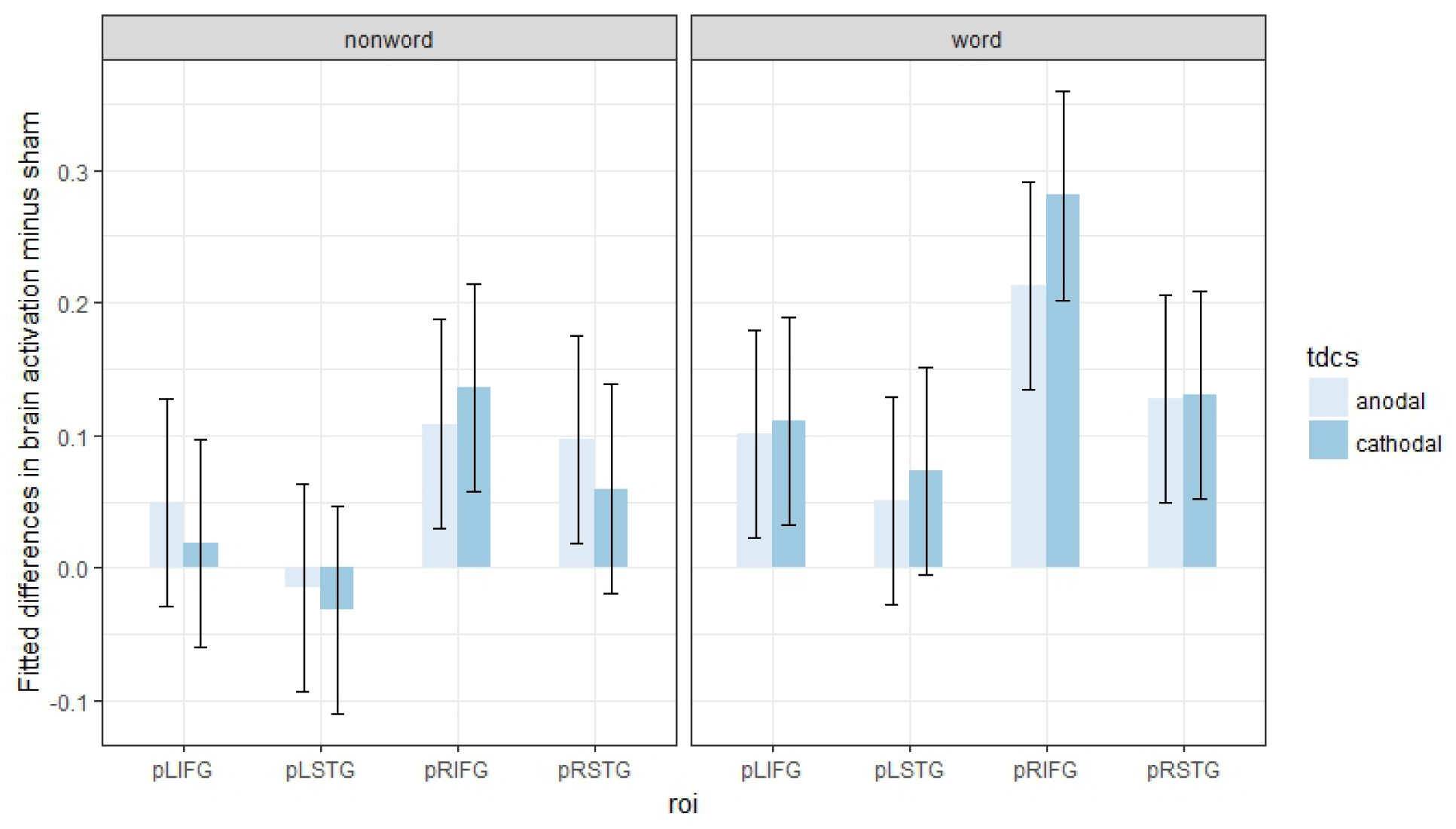
Fitted mean brain activation per ROI and stimulus type in lexical decision. The x-axis displays the ROIs. The y-axis displays the fitted mean brain activation. Error bars represent the contrast estimate ± the pooled standard error.

#### 3.4.2 Connectivity analysis per stimulus type and tDCS condition

Partial correlation analyses were performed between the fitted mean brain activations of the target network ROIs by stimulus type (i.e., word and nonword) and tDCS. Only anodal tDCS with the stimulus type word induced a significant correlation, the RIFG/LSTG (Tables 11 through 14).

### 3.5 Analysis of words and nonwords in word naming

#### 3.5.1 Stimulus type and tDCS effects on ROI mean brain activation

A 2 x 3 x 4 (stimulus type x tDCS x ROI) linear mixed effect model was fitted to the mean parameter estimates of ROI activation data of words and nonwords from the word naming task data. No significant interaction was observed.

Post hoc contrast analyses (Benjamini-Hochberg corrected for multiple comparisons) were conducted, but no significant result was observed (Figure 7 and Table 15).

**Table 11.** Partial correlation analyses for fitted mean brain activations in words of lexical decision under anodal tDCS

|  | LIFG | LSTG | RIFG | RSTG |
| --- | --- | --- | --- | --- |
| LIFG | 1.00 | 0.74 | -0.63 | 0.23 |
| LSTG | 0.74 | 1.00 | **0.96** | 0.47 |
| RIFG | -0.63 | 0.96 | 1.00 | -0.53 |
| RSTG | 0.23 | 0.47 | -0.53 | 1.00 |

Pearson's $r$ for each pair of ROIs with significant correlations at $p \leq 0.05$ marked in bold.

**Table 12.** Partial correlation analyses for fitted mean brain activations in nonwords of lexical decision under anodal tDCS

|  | LIFG | LSTG | RIFG | RSTG |
| --- | --- | --- | --- | --- |
| LIFG | 1.00 | -0.12 | 0.81 | 0.84 |
| LSTG | -0.12 | 1.00 | 0.21 | 0.40 |
| RIFG | 0.81 | 0.21 | 1.00 | -0.70 |
| RSTG | 0.84 | 0.40 | -0.70 | 1.00 |

Pearson's $r$ for each pair of ROIs. No significant correlation at $p \leq 0.05$ was observed.

**Table 13.**
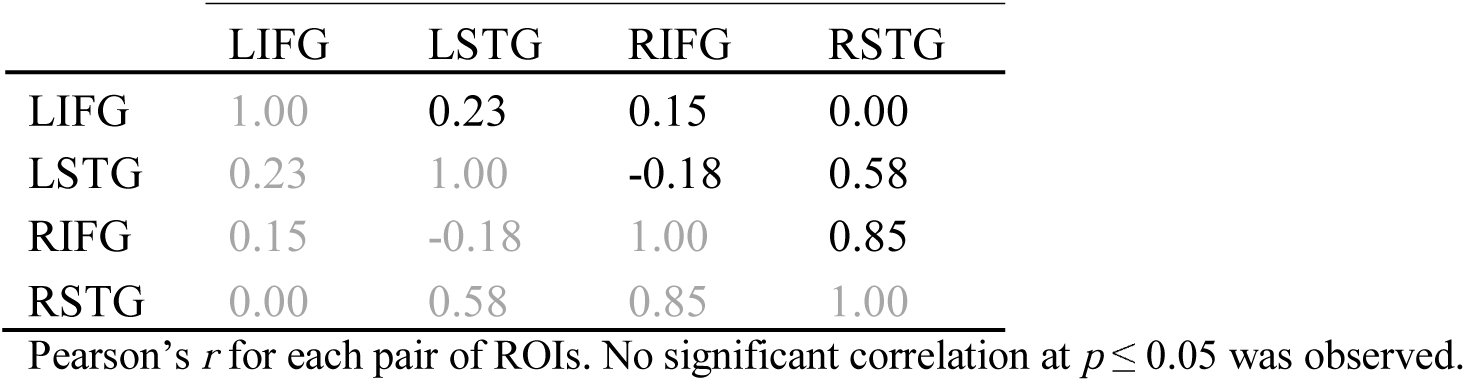
Partial correlation analyses for fitted mean brain activations in words of lexical decision under cathodal tDCS

**Table 14.**
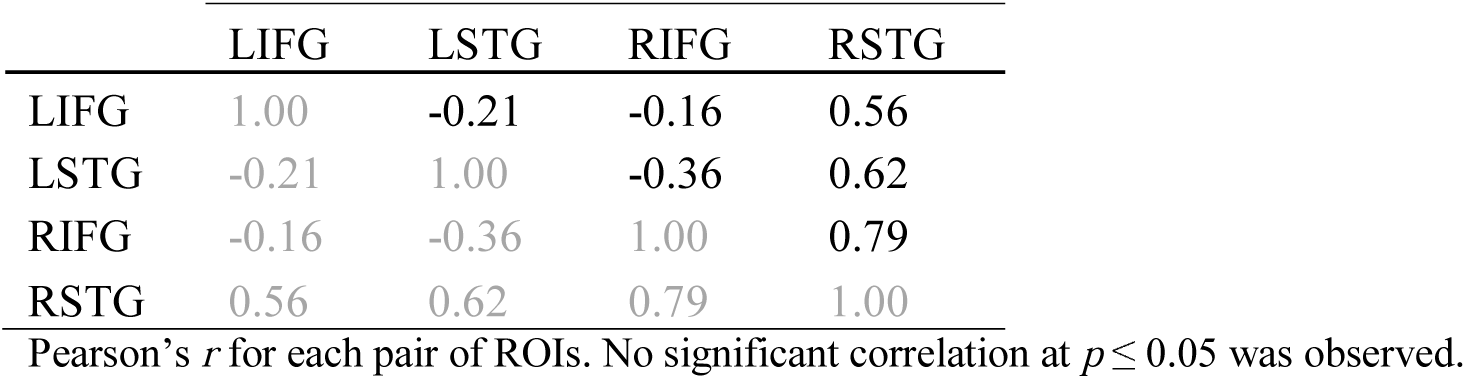
Partial correlation analyses for fitted mean brain activations in nonwords of lexical decision under cathodal tDCS

|  | LIFG | LSTG | RIFG | RSTG |
| --- | --- | --- | --- | --- |
| LIFG | 1.00 | -0.21 | -0.16 | 0.56 |
| LSTG | -0.21 | 1.00 | -0.36 | 0.62 |
| RIFG | -0.16 | -0.36 | 1.00 | 0.79 |
| RSTG | 0.56 | 0.62 | 0.79 | 1.00 |

Pearson's $r$ for each pair of ROIs. No significant correlation at $p \leq 0.05$ was observed.

**Table 15.** Contrast analyses for fitted brain activations per ROI and stimulus type

| Contrast | Estimate | SE | df | *t* | *p* |
| --- | --- | --- | --- | --- | --- |
| LIFG: nonword/anodal | 0.02 | 0.07 | 5 | 0.30 | 0.90 |
| LIFG: word/anodal | -0.06 | 0.07 | 5 | -0.78 | 0.90 |
| LIFG: nonword/cathodal | 0.08 | 0.07 | 5 | 1.01 | 0.90 |
| LIFG: word/cathodal | -0.01 | 0.07 | 5 | -0.11 | 0.98 |
| LSTG: nonword/anodal | 0.06 | 0.07 | 5 | 0.82 | 0.90 |
| LSTG: word/anodal | 0.03 | 0.07 | 5 | 0.42 | 0.90 |
| LSTG: nonword/cathodal | 0.06 | 0.07 | 5 | 0.75 | 0.90 |
| LSTG: word/cathodal | 0.02 | 0.07 | 5 | 0.29 | 0.90 |
| RIFG: nonword/anodal | 0.04 | 0.07 | 5 | 0.50 | 0.90 |
| RIFG: word/anodal | -0.08 | 0.07 | 5 | -1.09 | 0.90 |
| RIFG: nonword/cathodal | 0.08 | 0.07 | 5 | 1.04 | 0.90 |
| RIFG: word/cathodal | -0.04 | 0.07 | 5 | -0.60 | 0.90 |
| RSTG: nonword/anodal | 0.00 | 0.07 | 5 | -0.02 | 0.98 |
| RSTG: word/anodal | -0.05 | 0.07 | 5 | -0.72 | 0.90 |
| RSTG: nonword/cathodal | 0.09 | 0.07 | 5 | 1.23 | 0.90 |
| RSTG: word/cathodal | 0.04 | 0.07 | 5 | 0.48 | 0.90 |

**Figure 7.**
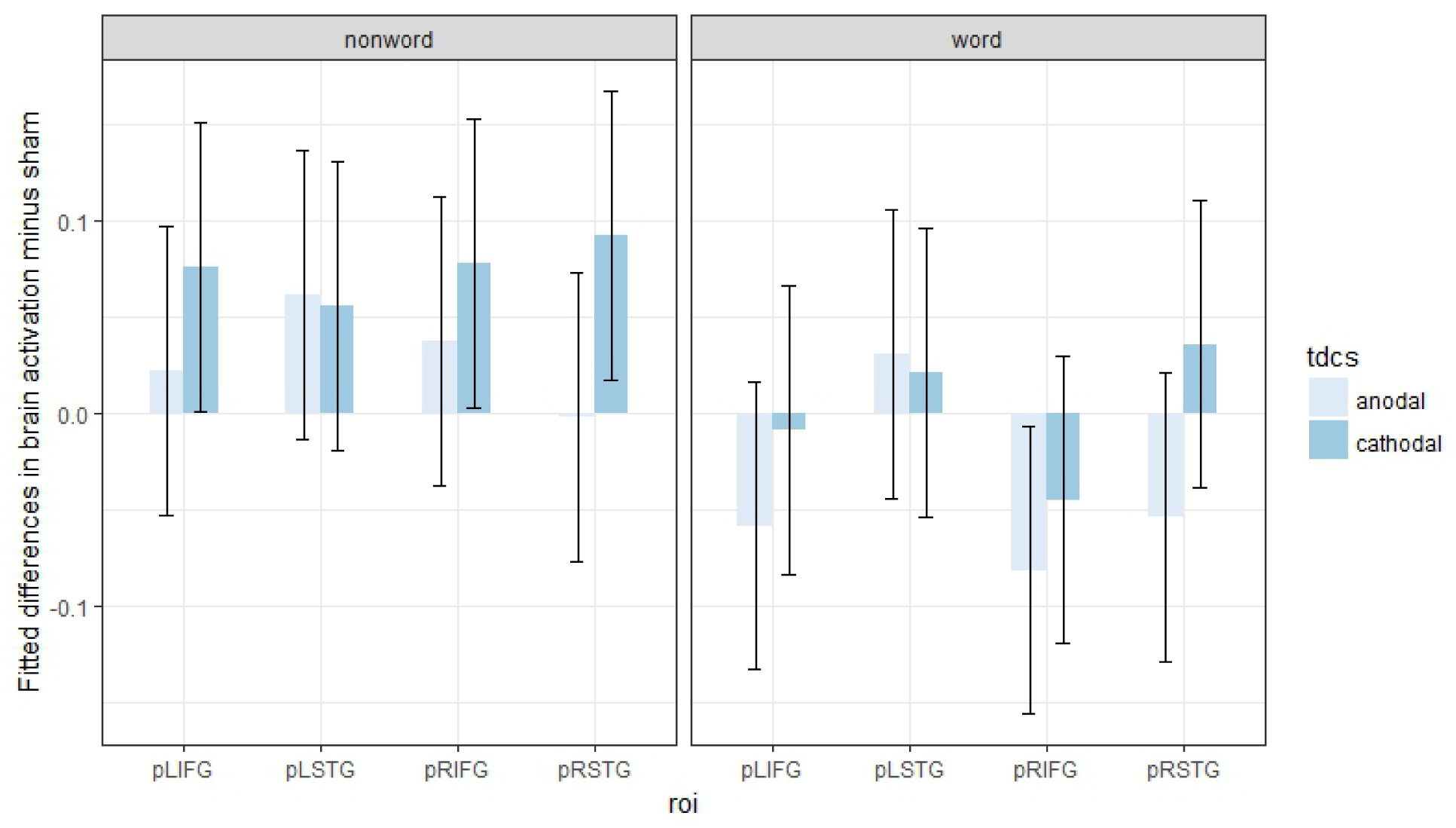
Fitted mean brain activation per ROI and stimulus type in word naming. The x-axis displays the ROIs. The y-axis displays the fitted mean brain activation. Error bars represent the contrast estimate ± the pooled standard error.

#### 3.5.2 Connectivity analysis per stimulus type and tDCS condition

Partial correlation analyses were performed between the fitted mean brain activations of the target network ROIs by stimulus type (i.e., word and nonword) and tDCS condition. Only anodal tDCS with stimulus type word induced significant correlations, which were the LIFG/RSTG, LIFG/LSTG, LIFG/RIFG, LSTG/RSTG and RIFG/RSTG (Tables 16 through 19).

**Table 16.** Partial correlation analyses for fitted mean brain activations in words of word naming under anodal tDCS

|  | LIFG | LSTG | RIFG | RSTG |
| --- | --- | --- | --- | --- |
| LIFG | 1.00 | **0.96** | **0.97** | **-0.97** |
| LSTG | 0.96 | 1.00 | -0.90 | **0.96** |
| RIFG | 0.97 | -0.90 | 1.00 | **0.97** |
| RSTG | -0.97 | 0.96 | 0.97 | 1.00 |

Pearson's *r* for each pair of ROIs with significant correlations at $p \leq 0.05$ marked in bold.

**Table 17.** Partial correlation analyses for fitted mean brain activations in nonwords of word naming under anodal tDCS

|  | LIFG | LSTG | RIFG | RSTG |
| --- | --- | --- | --- | --- |
| LIFG | 1.00 | -0.16 | 0.83 | -0.66 |
| LSTG | -0.16 | 1.00 | 0.53 | 0.05 |
| RIFG | 0.83 | 0.53 | 1.00 | 0.70 |
| RSTG | -0.66 | 0.05 | 0.70 | 1.00 |

Pearson's $r$ for each pair of ROIs. No significant correlation at $p \leq 0.05$ was observed.

**Table 18.**
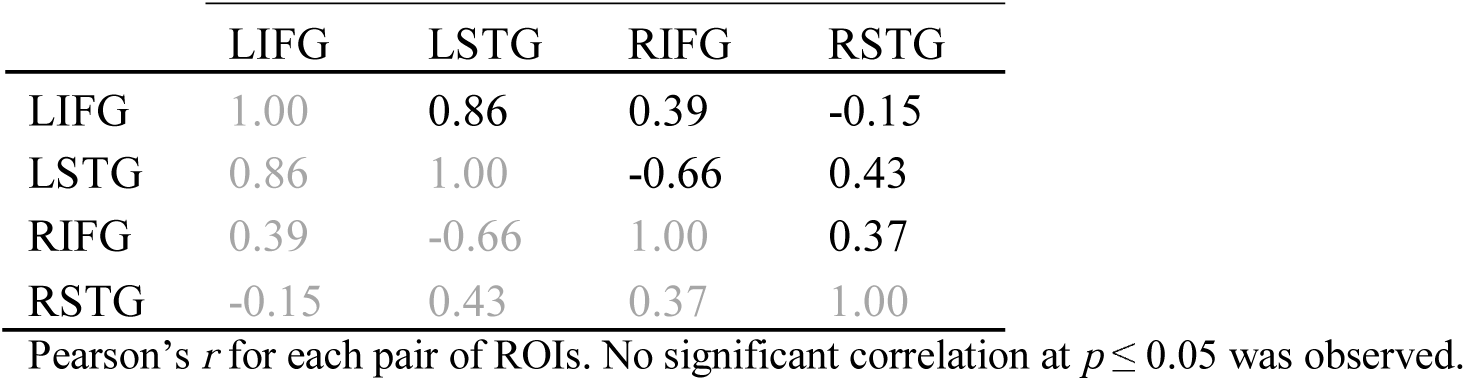
Partial correlation analyses for fitted mean brain activations in words of word naming under cathodal tDCS

|  | LIFG | LSTG | RIFG | RSTG |
| --- | --- | --- | --- | --- |
| LIFG | 1.00 | 0.86 | 0.39 | -0.15 |
| LSTG | 0.86 | 1.00 | -0.66 | 0.43 |
| RIFG | 0.39 | -0.66 | 1.00 | 0.37 |
| RSTG | -0.15 | 0.43 | 0.37 | 1.00 |

Pearson's $r$ for each pair of ROIs. No significant correlation at $p \leq 0.05$ was observed.

**Table 19.** Partial correlation analyses for fitted mean brain activations in nonwords of word naming under cathodal tDCS

|  | LIFG | LSTG | RIFG | RSTG |
| --- | --- | --- | --- | --- |
| LIFG | 1.00 | 0.81 | -0.17 | 0.35 |
| LSTG | 0.81 | 1.00 | 0.27 | 0.15 |
| RIFG | -0.17 | 0.27 | 1.00 | 0.17 |
| RSTG | 0.35 | 0.15 | 0.17 | 1.00 |

Pearson's $r$ for each pair of ROIs. No significant correlation at $p \leq 0.05$ was observed.

## 4. DISCUSSION

The aim of this study was to investigate how the dyslexic brain would handle tDCS perturbations during the performance in tasks which involve phonological processing, a language function that is affected in PWD (Brunswick et al., 1999; Ramus et al., 2004). More specifically, we investigated whether task load of tasks involving phonological processing would have a differential effect on the outcomes of tDCS stimulation of the LIFG in PWD. To accomplish this aim, tasks covering a range from speech perception to speech production were used. This range was assumed to pose increasingly higher task load to the LIFG (Amunts et al., 1999; Eickhoff et al., 2009; Indefrey, 2011; Lee et al., 2012; Liakakis et al., 2011; Liebenthal et al., 2013).

Typical predictions for the effects of tDCS as a function of task load were as follows for the tDCS of the LIFG. Anodal tDCS was expected to increasingly induce facilitation across the speech perception to speech production range of tasks, whilst cathodal stimulation was expected to increasingly induce inhibition. Cathodal tDCS was also expected to induce facilitation via compensation, which was more likely to occur in conditions of lower task load. However, due to the phonological processing deficit in PWD, some of these typical predictions could not apply or weaker responses (for example in terms of brain activation or number of significant connections between network nodes) could be generated for PWD when compared to predictions for healthy young adults. For example, compensation of the downregulation of the LIFG for a task of speech perception was expected to be at least weaker. This is because the LSTG, a brain area highly relevant for phonological processing in speech perception tasks (Lee et al., 2012; Liebenthal et al., 2013), is known to be hypoactivated in PWD (Brunswick et al., 1999; Georgiewa et al., 1999; Hoeft et al., 2006; Ruff et al., 2003), and therefore less likely to offer compensation. Results of this study suggest that the baseline brain activation associated with the phonological deficit in PWD restricts the network strategies that can be used by the brain and modulates responses that otherwise should follow predictions based on task load for the healthy brain.

Outcomes were evaluated in terms of significant prominent connections between the target network nodes induced by the direct current. In general, results of both anodal tDCS and cathodal tDCS were contrary to predictions based on task load for the healthy brain. Anodal tDCS of the LIFG should increase facilitation, whilst cathodal tDCS should increase inhibition, across the tasks in the range from speech perception to speech production and from words to nonwords. Compensation induced by cathodal tDCS was expected to be decreasingly lower. Contrary to task load based predictions, anodal tDCS caused larger facilitation for categorical perception than for word naming, whilst cathodal tDCS-induced compensation was larger for word naming than for categorical perception. Similarly, in both the lexical decision and the word naming tasks, anodal tDCS facilitation was larger for words than for nonwords, whereas cathodal tDCS had an apparently equivalent inhibitory effect for both stimulus types, when compensation was expected for words. It should be noted, however, that in all these comparisons, the part said to have smaller facilitatory effect had non-significant results, that were interpreted as so for theoretical reasons. Nevertheless, since no definitive conclusions can be drawn from non-significant results, this interpretation should be taken cautiously.

Taken together, these brain stimulation results and the known pattern of inefficient brain activation (LIFG hyperactivation compensating for the LSTG hypoactivation) for phonological processing in PWD suggest a (maladaptive) shift of function from the LSTG to the LIFG (Brunswick et al., 1999). Consequently, the roles previously attributed to the LIFG cannot be performed at a suitable level. This interpretation would explain findings where the LIFG showed to be more responsive to anodal tDCS during the categorical perception task and with the stimulus type words (for both lexical decision and word naming) instead of during the word naming task and with the stimulus type nonwords, as justified by the new LIFG duties. Because the LIFG is overloaded and its multi-task role seems maladaptive, the inconsistent pattern of responses to old and new duties seems also justified. For example, cathodal tDCS strongly downregulated the LIFG (no significant prominent connections; as discussed above, a theoretically motivated tentative interpretation of a null finding) for nonwords in both lexical decision and word naming. Since no sign of compensation was observed, this indicates that nonwords had a high task load for the LIFG, as assumed for the healthy brain.

## Acknowledgements

This work was supported by a postgraduate scholarship from Coordenação de Aperfeiçoamento de Pessoal de Nível Superior (CAPES) (Brazil)/University of Birmingham to L.R.A.

